# Equivalence of charge imbalance and external electric fields during free energy calculations of membrane electroporation

**DOI:** 10.1101/2023.01.13.523896

**Authors:** Gari Kasparyan, Jochen S. Hub

**Affiliations:** Theoretical Physics and Center for Biophysics, Saarland University, Saarbrücken, Germany

## Abstract

Electric fields across lipid membranes play important roles in physiology, medicine, and biotechnology, rationalizing the wide interest in modeling transmembrane potentials in molecular dynamics simulations. Transmembrane potentials have been implemented with external electric fields or by imposing charge imbalance between the two water compartments of a stacked double-membrane system. We compare the two methods in the context of membrane electroporation, which involves a large change of membrane structure and capacitance. We show that, given that Ewald electrostatics are defined with tinfoil boundary conditions, the two methods lead to (i) identical potentials of means force (PMFs) of pore formation and expansion at various potentials, demonstrating that the two methods impose equivalent driving forces for large-scale transitions at membranes and (ii) to identical polarization of water within thin water wires or open pores, suggesting that the two methods furthermore impose equivalent local electric fields. Without tinfoil boundary conditions, effects from external fields on pore formation are spuriously suppressed or even removed. Together, our study shows that both methods, external fields and charge imbalance, are well suitable for studying large-scale transitions of lipid membranes that involve changes of membrane capacitance. However, using charge imbalance is technically more challenging for maintaining a constant transmembrane potential since it requires updating of the charges as the capacitance of the membrane changes.

## Introduction

Electric fields over lipid membranes play key roles in physiology and biotechnology, explaining the wide interest in modeling transmembrane potentials in molecular dynamics (MD) simulations. Transmembrane potentials have been applied in a plethora of MD studies involving, among others, permeation across ion channels^1–3^ or water channels,^4^ voltage gating,^5,6^ or membrane electroporation.^7–12^ These simulations provided unprecedented mechanistic and energetic insight into complex transitions at biological membranes. Membrane electroporation, which is in the focus of the present study, denotes the formation of pores into lipid bilayers by the application of transmembrane electric potentials.^13–15^ Reversible electroporation is widely used in biotechnology or medicine to deliver biological samples such as genes, vaccines, or drugs into living cells. ^16^ Furthermore, irreversible electroporation is used to kill malignant cells for the ablation of tumor tissue that is not accessible to surgery.^17,18^

Two methods have been put forward for implementing transmembrane potentials in MD simulations. First, external electric fields **E** have been applied along the membrane normal by applying an additional force *q_i_***E** to the atoms, where *q_i_* denotes the partial charge of atom *i*.^19–21^ With an external electric field with magnitude *E_z_* along the *z*-axis of the simulation box, the overall potential drop along the box is given by *E_z_L_z_*, where *L_z_* is the box dimension along the *z* direction.^21,22^ In presence of a lipid membrane oriented in the *x-y* plane, the potential drops mostly over the low-dielectric membrane core, hence imposing a transmembrane potential of *V_m_* = *E_z_L_z_*. Several studies used external electric fields to drive membrane electroporation.^7,8,10–12,23,24^

Alternatively, transmembrane potentials have been implemented by simulating a system with two stacked membranes, thereby forming two solvent reservoirs.^4,9,12,25,26^ In this setup, the potential is imposed using a charge imbalance between the reservoirs, i.e., by adding ions with excess charges +*Q* and –*Q* into the two solvent reservoirs, respectively. This setup has been referred to as “charge imbalance”, “ion imbalance” or “double-membrane setup”. The charge imbalance *Q* required for obtaining a specific transmembrane potential *V_m_* is not obvious since it depends in the capacitance *C* of the membrane via *Q* = *CV_m_*, while the membrane capacitance depends on the thickness, internal structure, and protein content. Hence, *Q* is typically modified by trial and error until the observed transmembrane potential agrees with the desired value. Upon structural transitions of the membrane such as the formation of a transmembrane pore, the capacitance may greatly change, suggesting that *Q* must be updated for maintaining a constant *V_m_*. An extension of the charge imbalance setup is given by the “computational electrophysiology” method, which maintains a constant transmembrane potential after ion permeation by placing back permeated ions.^27^

De-facto standards of biomolecular MD simulations involve the use of three-dimensional periodic boundary conditions to avoid surface artifacts and the use of Ewald sums^28^ for evaluating the long-range electrostatic interactions. Many modern MD codes implement Ewald sums via the computationally efficient particle-mesh Ewald (PME) method.^29,30^ Ewald sums are typically evaluated assuming so-called tinfoil boundary conditions, implying that the periodic lattice of unit cells is at a large distance surrounded by a conducting medium. However, the use of Ewald sums with tinfoil boundary conditions together with external fields has been considered as problematic because the external field induces a macroscopic dipole in the simulation box, which seems being at odds with the assumption of a conducting surrounding medium. ^31^ Hence, it has been discussed whether the external field may be straightforwardly compared with fields under experimental conditions.^10,11,31,32^ In a similar spirit, spurious water ordering effects due to the presence of a macroscopic dipole in simulation systems with asymmetric lipid membrane have been described recently.^33^

Several studies compared the effects of external electric fields with effects of charge imbalance.^12,34^ Melcr *et al*. showed that the two methods yield equivalent ion and charge density distributions, water dipole orientations, and local electric fields around a lipid bilayer. However, since Melcr *et al*. focused on planar membranes, it has not been tested whether the two methods are equivalent during transmembrane pore formation, which involves large changes of membrane capacitance. In addition, is remained unclear whether electric fields implemented with the two methods impose equivalent free energy gradients for conformational transitions that are driven by transmembrane potentials.

Here, we simulated membrane electroporation across a dipalmitoylphosphatidylcholine (DPPC) membrane using either external electric fields or the charge imbalance method. As a highly sensitive test for the effect of electric fields on the membrane, we computed the potential of mean force (PMF) for pore nucleation and expansion at transmembrane potentials between 0 mV and 600 mV using a recently proposed reaction coordinate.^35^ As a probe for the local electric fields acting in the simulation, we computed the average water dipole orientation along the pore axis. We find that, given that PME is applied with tinfoil boundary conditions, external electric fields and charge imbalance yield nearly identical PMFs and dipole orientations, suggesting that the two methods are equally suitable for studying large-scale transitions at biomembranes.

## Methods

### Simulation setup and parameters

The simulation system of a single membrane, composed of 200 dipalmitoylphosphatidylcholine (DPPC) lipids and 12000 water molecules, was set up with the MemGen web server.^36^ Lipid interactions were described with the force field by Berger *et al*.,^37^ and the SPC water model was applied.^38^ The system was equilibrated with the GROMACS simulation software, version 2018.^39^ Electrostatic interactions were described with the PME method with tinfoil boundary conditions if not stated otherwise, implying that the periodic simulation system is surrounded by a conducting medium.^29,30^ Dispersion interactions and short-range repulsion was described by a Lennard-Jones potential with a cutoff at 1 nm. The temperature was kept at 323 K using velocity-rescaling (*τ* = 0.1 ps).^40^ The pressure was controlled at 1 bar using weak coupling (*τ* = 5 ps).^38^ A time step of 5 fs was applied. The system was equilibrated until the potential energies and box dimensions were fully converged.

### Reaction coordinate of pore formation

PMFs were computed along a recently proposed joint reaction coordinate *ξ*_p_ for pore nucleation and pore expansion.^35^ During pore nucleation, *ξ*_p_ is equivalent to the chain reaction coordinate *ξ*_ch_ that quantifies the degree of connectivity of a polar transmembrane defect and takes values *ξ*_ch_ ∈ [0, 1).^41,42^ It was previously shown that PMF calculations along *ξ*_ch_ using umbrella sampling (US) simulations do not suffer from hysteresis problems and yield converged PMFs within moderate simulation times, both in the context of pore formation^42–44^ or stalk formation.^45^ The coordinate *ξ*_p_ used in this study has been defined as^35^

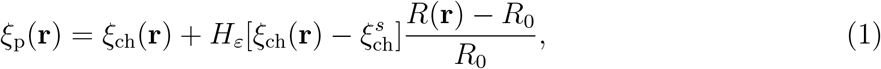

where *H_ε_* is a differentiable variant of the Heaviside step function with a switch interval [–*ε, ε*], *R*(**r**) is the radius of an established transmembrane pore, *R*_0_ = 0.419 is the approximate radius of a thin water defect, and 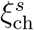 is the position of *ξ*_ch_ of a thin water defect. Hence, for 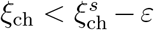, the coordinate *ξ*_p_ is equivalent to *ξ*_ch_ and quantifies the connectivity of the transmembrane defect. For 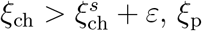 is given by the pore radius *R*(**r**) in units of *R*_0_. For more details on the definition of *ξ*_ch_(**r**) and *R*(**r**) we refer to previous work.^35,41^

Additional parameters required for defining of *ξ*_p_ were chosen as follows. The polar atoms contributing to *ξ*_p_ were taken as the oxygen atoms of water as well as the four oxygen atoms lipid phosphate groups. The switch region between pore nucleation and pore expansion was defined with the parameters 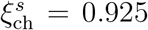 and *ε* = 0.05. The radius *R*(**r**) of the open pore was computed from the number of polar atoms within a horizontal layer centered in the membrane plane with the thickness of 1.2 nm, where the volume per polar atom was taken as 0.02996 nm^3^ corresponding to the volume per water molecule. The cylinder used for defining the chain coordinate *ξ*_ch_ was composed of 28 slices with a thickness of 0.1 nm each, and using a cylinder radius of *R_cyl_* = 1.2 nm. Critically, during pore nucleation, the radius of the defect is fully controlled by force field (together with other simulation parameters such as the temperature) and not by the cylinder radius, as the latter is purely used to control the locality of the defect in the membrane plane. The cylinder ensures that two laterally displaced partial defects in the membrane connected with the upper or lower solvent reservoir, respectively, are not misinterpreted as a continuous transmembrane defect, which would lead to hysteresis problems. The length of 2.8 nm of the cylinder was chosen such that approximately 25% of the cylinder slices were filled by polar atoms (*ξ*_ch_ ≈ 0.25) in the flat membrane.

### Umbrella sampling simulations of pore formation

Pores were formed by constant-velocity pulling simulations along *ξ*_p_ over 100 ns from *ξ*_p_ = 0 to *ξ*_p_ = 5 using a force constant of 3000 kJ mol^-1^. Initial frames for umbrella sampling (US) simulations were taken from the constant-velocity pulling simulation. The PMFs were computed using 52 umbrella windows. Since sampling is more challenging in the switch region between pore nucleation and expansion (i.e., where 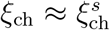), we used the following non-equidistantly distributed US reference positions and non-equal US force constants: 0.08 to 0.64 in steps of 0.08 (force constant *k* = 3000 kJ mol^-1^); 0.69 to 1.03 in steps of 0.02 (*k* = 5000 kJ mol^-1^); and 1.18 to 4.93 in steps of 0.15 (*k* = 400 kJ mol^-1^). Each window was simulated for 100 ns, and the first 40 ns were removed for equilibration. Statistical errors were estimated using 50 rounds of the Bayesian bootstrap of complete histograms.^46^

### Charge imbalance (CI) simulations

For the CI simulations, a pre-equilibrated simulation system of 200 DPPC lipids (see above) was doubled along the *z* direction (membrane normal) with the GROMACS module gmx genconf. A grid of 4 × 4 dummy charges was placed at the center of each of the two solvent compartments (Fig. 1B, blue and red spheres). The positions of the dummy charges were restrained with harmonic potentials with force constant 10 kJ mol^-1^nm^-2^. The Lennard-Jones parameters of the dummy charges were taken from the potassium ion of the GROMOS force field^47^ as C_6_ = 1.63787 × 10^-5^ kJ mol^-1^ nm^6^ and C_12_ = 1.18384 × 10^-6^ kJ mol^-1^nm^12^.

**Figure 1:**
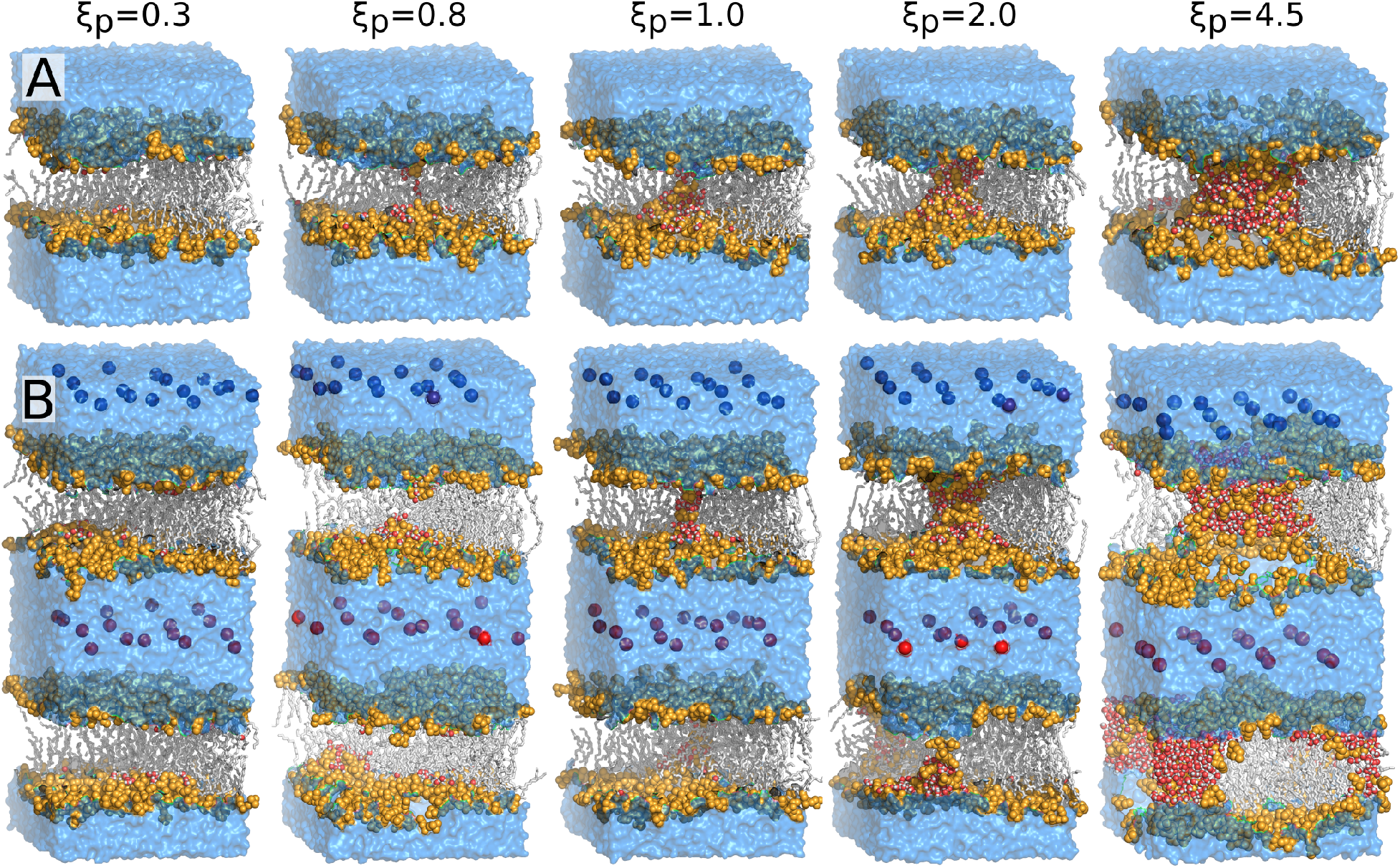
Typical simulations snapshots of pore nucleation and pore expansion, shown for (A) the single-membrane system with external electric field and (B) for the double-membrane system with charge imbalance. Corresponding values of the reaction coordinate *ξ*_p_ are shown, which quantifies the degree of connectivity of the transmembrane defect for *ξ*_p_ ≲ 1 and the radius R of the pore via R = *ξ*_p_*R*_0_ during pore expansion for *ξ*_p_ ≳ 1 (*R*_0_ = 0.419 nm). Headgroup atoms are shown as orange spheres, lipid tails as grey sticks, water in the pore as red/white spheres, and other water as blue surface. Restrained positive and negative dummy charges used to impose the charge imbalance are shown as blue and red spheres, respectively.

In order to define dummy charges that yield a desired transmembrane potential, a series of equilibrium simulations was carried out: 20 ns with dummy atoms with charge set to 0e and another 20 ns using charges of 0.1e, where e denotes the unit charge. The resulting transmembrane potential was computed using the gmx potential module, which derives the electrostatic potential profile *ϕ*(*z*) via a double integration of the one-dimensional Poisson equation. Here, the potential was taken as the difference between the two flat segments of *φ*(*z*) corresponding to the two water regions. According to these simulations, we found that, for an intact membrane, the imposed transmembrane potential followed approximately *V_m_* = *Q* · 5.7 V/C, where *Q* is the total absolute charge of each of the charge grids. From this analysis, the initial charges were used during US simulations of pore opening.

### Iterative optimization of charge imbalance over rounds of umbrella sampling

The capacitance *C* of the membrane increases with increasing pore radius, suggesting that the charge imbalance *Q* must be increased in order to maintain a constant transmembrane potential *V_m_* = *Q/C*. We optimized the charge imbalance *Q* for each umbrella window in an iterative manner. After each round of US, we computed the transmembrane potential and updated *Q* to obtain increasingly better agreement with the target potential. To reduce the effects of statistical fluctuations, we smoothed the computed potentials along neighboring umbrella windows using a moving average with a window size of three. If the smoothed potential for an umbrella window 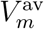 differed from the target potential 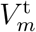 by more than 10%, we scaled the charges by 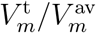. If the relative deviation was smaller than 10%, we considered the charges as converged.

## Results and Discussion

Figure 1 presents typical simulations frames along the pore nucleation and pore expansion pathway, taken from final frames of US windows. Frames are shown for the single-membrane system subject to an external electric field (Fig. 1A) and for the double-membrane system subject to a charge imbalance between the two water compartments (Fig. 1B). Pore nucleation proceeds from the flat membrane (*ξ*_p_ = 0.25) via thinning of the membrane involving the protrusion of a polar defect into the membrane core (*ξ*_p_ = 0.85), the formation of a highly transient water needle, up to the formation of a membrane-spanning polar defect (*ξ*_p_ = 1). As the pore forms, lipids reorient along the pore rim to partly shield the hydrophobic membrane core from the aqueous defect, as visible from the lipid head groups along pore rim (Figs. 1A/B, right panels). Hence, lipid and water conformations observed during US simulations agree with a large number of previous simulations of pore formation^7,8,43,48–52^

### Maintaining a constant transmembrane potential in the double-membrane systems requires updating of the charge imbalance

Figure 2A shows typical potential profiles Δ*V*(*z*) for the double membrane system for trans-membrane potentials *V_m_* between 0mV and 600 mV, here taken from an US simulation with a pore of radius ~1.25nm (*ξ*_p_ = 3). The two flat regions at |*z*| ≈ 9 nm and |*z*| ≈ 0 nm correspond to the two solvent reservoirs that embed the dummy charge layers used to impose the potential. The potential largely drops over the hydrophobic cores of two membranes (3nm < |*z*| < 6nm), as expected due to the lower dielectric permittivity of the membrane cores as compared to the permittivity of the water compartments. The transmembrane potential *V_m_* imposed by the dummy charges is given the potential difference between the two solvent reservoirs.

**Figure 2:**
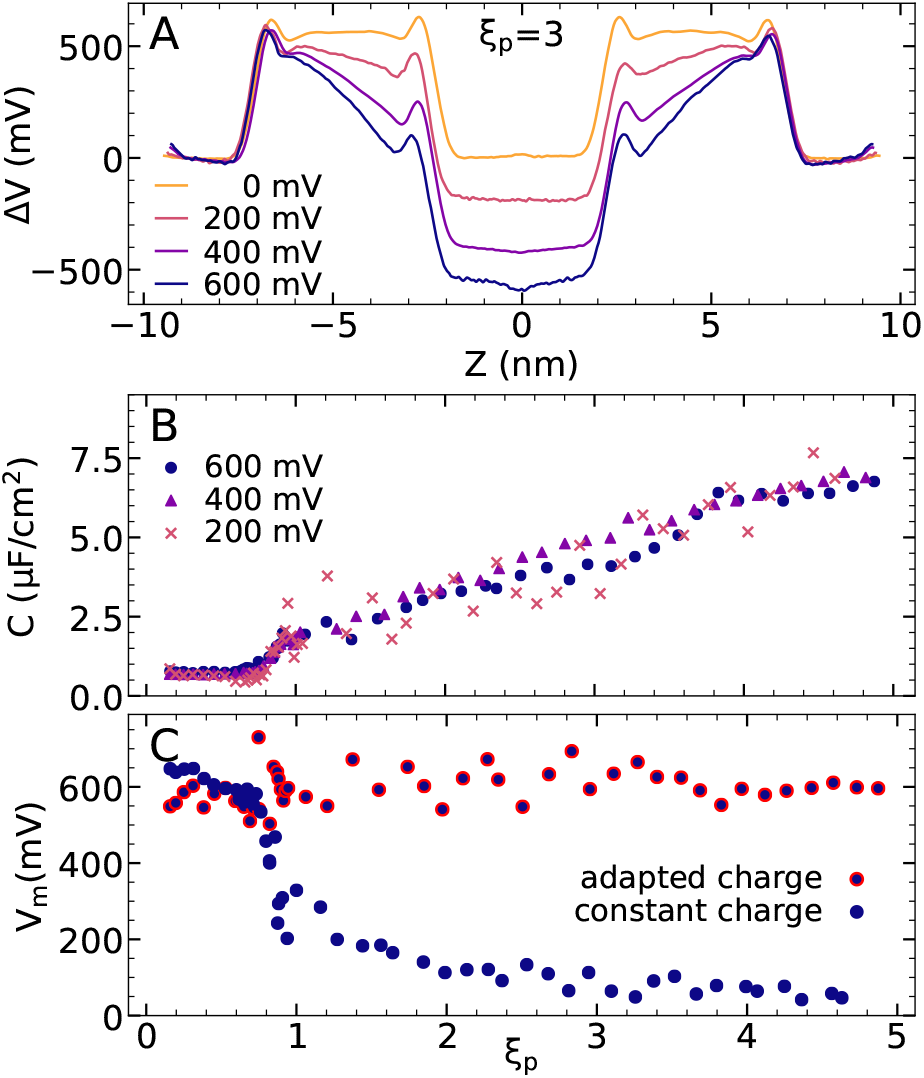
(A) Electrostatic potential profiles Δ*V*(*z*) along the membrane normal z in double membrane systems with potentials of 0mV, 200mV, 400mV, and 600mV (see legend). (B) Capacitance *C* of the membrane versus the reaction coordinate of pore formation and pore expansion *ξ*_p_, computed from the transmembrane potential *V_m_* and the charge imbalance *Q* via *C* = *Q/V_m_*. Symbols (see legend) correspond to values computed from individual US windows. (C) Transmembrane potential obtained from the Δ*V*(*z*) profiles (cf. panel A) with a target potential 600 mV, computed from US windows of various *ξ*_p_ after adapting the charge imbalance for maintaining a constant *V_m_* (red circles) or using a constant charge imbalance (blue circles).

As the pore expands, the capacitance of the membrane increases because part of the low-dielectric membrane core is replaced with high-dielectric water (Fig. 2B). Consequently, in the double-membrane system, using a constant charge imbalance would lead to a drop of *V_m_* (Fig. 2C, blue circles). To maintain a constant, pre-selected potential *V_m_* during pore expansion requires updating of the dummy charges. We found that *V_m_* converges slowly within individual US windows, possibly owing to slow rearrangements of the pore shape and of the lipid headgroup structure along the pore rim on the time scale of several tens of nanoseconds. The slow convergence of *V_m_* complicates the identification of the correct charge imbalance that leads to a pre-selected *V_m_*. Nevertheless, using an iterative procedure, we identified charge imbalances that kept the *V_m_* reasonably constant over all US windows, as shown in Fig. 2C (red circles) for simulations with 600 mV.

### PMFs of electroporation agree between charge imbalance and external field simulations

We computed the PMF of pore nucleation and pore expansion using either the double-membrane system (Fig. 3, blue lines) or using external electric fields (Fig. 3, orange lines). For the double-membrane system, two pores were formed simultaneously in the two membranes (Fig. 1B), and the US histograms from the two pore opening processes were combined into a single PMF, thereby providing the free energy per pore. In the PMFs shown in Fig. 3, the local minima at *ξ*_p_ ≈ 0.25 corresponding to the flat membrane were taken as reference point where the free energy was set to zero. The 0.25 ≤ *ξ*_p_ ≲ 0.9 range of the PMFs describes the nucleation of pores, whereas the *ξ*_p_ ≳ 1 region describes the expansion of fully formed polar defects with radii of approximately *R* = *ξ*_p_*R*_0_ with *R*_0_ = 0.419 nm. In qualitative agreement with previous simulations of spontaneous pore formation under non-equilibrium conditions^7,8,43,48–52^ and with theories of electroporation,^53^ application of electric fields leads to a large stabilization of the pore in a voltage- and radius-dependent manner. The physical implications of the PMFs will be discussed elsewhere. As a key finding of this study, Fig. 3 demonstrates that the PMFs based on charge imbalance reveal excellent agreement with the PMFs based on external electric fields. Considering that the PMFs are highly sensitive to modulations of *V_m_*, this agreement suggests that the two methods impose similar electrostatic environments, even during large-scale conformational transitions of membranes as studied here.

**Figure 3:**
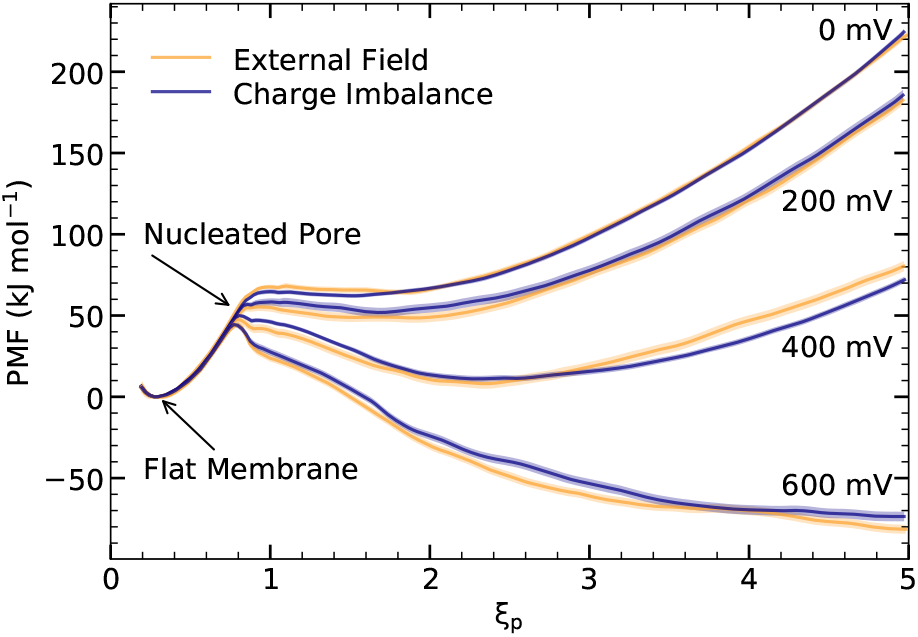
PMF of pore formation over a DPPC membrane involving pore nucleation (0 < *ξ*_p_ ≲ 0.9) and pore expansion (*ξ*_p_ ≳ 1) for potentials between 0mV, and 600 mV (see labels) using external electric fields (orange) or charge imbalance (blue).

### Analysis of water dipoles suggests identical electric fields in charge imbalance and external field simulations

As a probe for the local electric fields along the defect, we computed the average z component of water dipoles 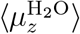 as function of z coordinate. In contrast to a similar analysis presented by Melcr et al.^34^ who studied the water dipoles for intact planar membrane, we here computed the dipoles for different degrees of pore opening by averaging within various US simulations. Figure 4 presents the 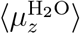 profiles taken from US windows with a thinned membrane (*ξ*_p_ = 0.7, yellow), with a thin defect with a radius of ~0.42nm (*ξ*_p_ = 1, orange), or with increasingly larger pores (*ξ*_p_ = 1.4, red; *ξ*_p_ = 3.2, purple; *ξ*_p_ = 4.9, blue). At the headgroup regions (|*z*| ≈ 2 nm), the profiles reveal preferred orientations of water dipoles that diminish with increasing pore opening. At the membrane center (|*z*| ≈ 0 nm), water in thin transmembrane defects are strongly polarized (Fig. 4B–D, *ξ*_p_ = 1 or *ξ*_p_ = 1.4, orange or red), rationalized by the fact that the membrane-crossing electric field lines concentrate within the higher dielectric. As the pore expands, the electric field decreases inside the pore, leading to decreasingly oriented water dipoles (Fig. 4B–D, *ξ*_p_ ≥ 3.2, purple or blue).

**Figure 4:**
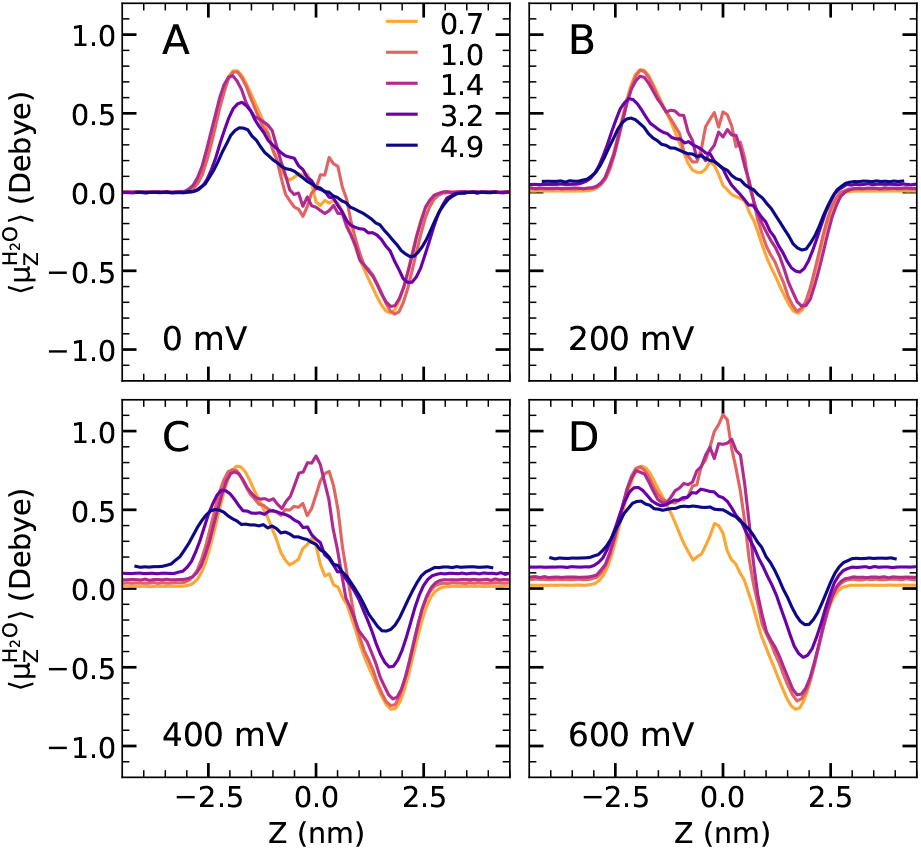
Average dipole per water molecule projected onto the z axis (membrane normal) for (A) 0mV, (B) 200mV, (C) 400mV, and (D) 600mV for increasing degrees of pore opening, as given by the *ξ*_p_ reaction coordinate at 0.7, 1.0, 1.4, 3.2, and 4.9 colored in shades from yellow to blue (see legend).

To test whether the local electric fields inside the pores agree between the double-membrane and the external field simulations, we compared 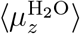 obtained from the two simulation setups. Figure 5 presents the profiles for systems with a thin pore with a radius of ~0.42nm (*ξ*_p_ = 1, solid lines) or with a wide open pore of radius ~2nm (*ξ*_p_ = 4.9, dashed lines). For systems with identical *V_m_* and identical degree of pore opening, we find excellent agreement between the charge imbalance (Fig. 5, yellow lines) and the external field systems (Fig. 5, blue lines). These data corroborate the conclusions from the PMFs that charge imbalance and external fields yield highly similar electrostatic environments across the membrane at different degrees of pore opening.

**Figure 5:**
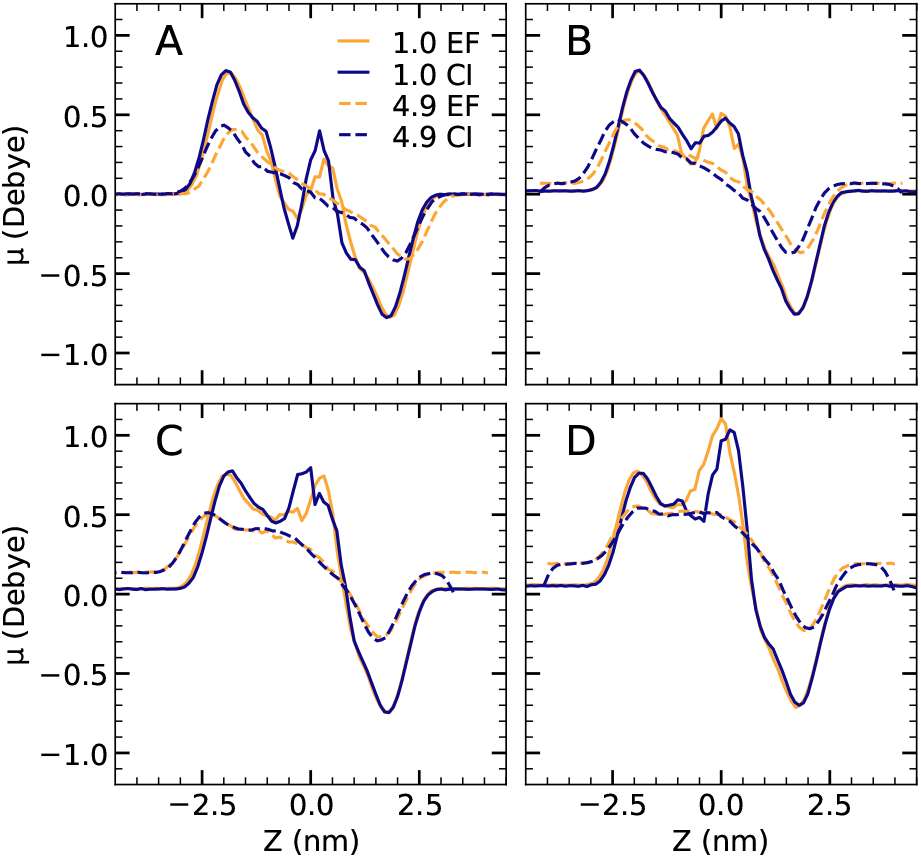
Profiles of average water dipoles, projected onto the z axis, for (A) 0mV, (B) 200 mV, (C) 400 mV, and (D) 600 mV, using the double-membrane system with charge imbalance (CI, blue) or the external electric field (EF, yellow). Profiles are shown for a thin defect (*ξ*_p_ = 1, solid lines) and for a wide open pore (*ξ*_p_ = 4.9, dashed lines). Excellent agreement between CI and EF simulations is found.

### Realistic external field simulations require tinfoil boundary conditions

According to the Ewald formula, the electrostatic energy per simulation unit cell can be decomposed into four contributions: the real space energy, the reciprocal space energy, the self-energy, and a dipole correction.^54,55^ The formula assumes that the periodic lattice of unit cells is at a large distance surrounded by a spherical boundary to a surrounding medium with dielectric permittivity *ε′*. The dipole correction is ^54,55^

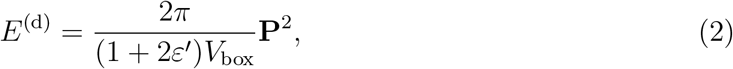

where **P** = Σ*_i_q_i_***r***_i_* is the macroscopic dipole of the system with atomic charges qi and positions ri, while *V*_box_ is the box volume. Biomolecular simulations are typically carried out with tinfoil boundary conditions (*ε′* = ∞) leading to a vanishing dipole correction. With finite *ε′*, in contrast, the dipole correction suppresses the formation of a macroscopic dipole.

To test the effect of the dipole correction during simulations with an external field, we repeated the PMF calculation of pore formation with *ε′* = 1 or *ε′* = 80, thereby modeling a surrounding vacuum or a surrounding aqueous medium, respectively (Fig. 6). In these simulations, we applied the external field that, with tinfoil boundary conditions, successfully modeled a transmembrane potential of 600 mV (cf. Fig. 3). According to the PMFs, using *ε′* = 80 instead of *ε′* = ∞ reduced the effect of the external field on the free energy of a large pore by ~50 kJ/mol (Fig. 6, compare red and orange curves). Using *ε′* = 1, the PMF with an external field (Fig. 6, purple) is nearly identical to the PMF computed without external field (Fig. 6, black), demonstrating that vacuum boundary conditions suppress the pore-stabilizing effect by the external electric field nearly completely.

**Figure 6:**
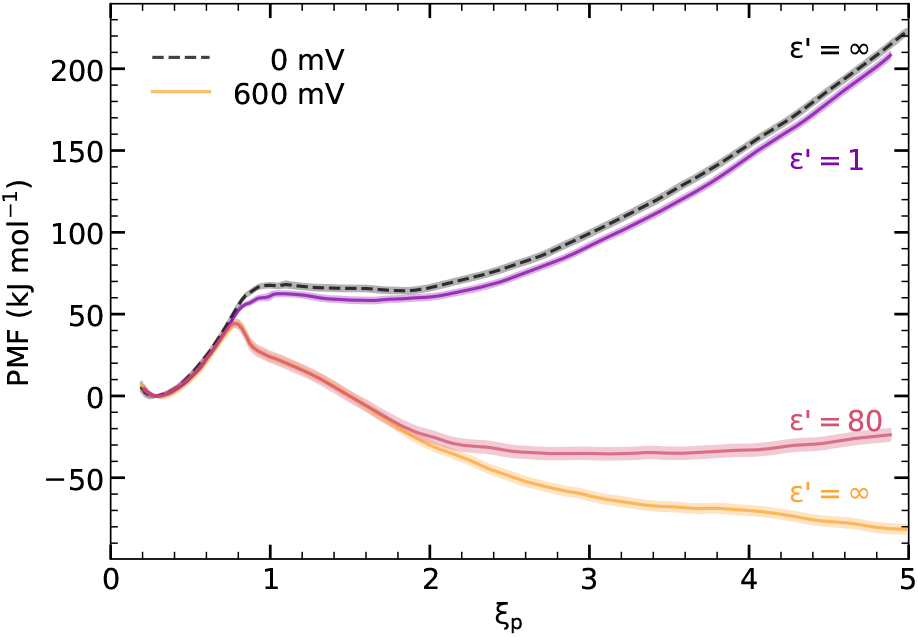
On the importance of using tinfoil boundary conditions: PMFs of pore formation using an external field of 0mV (black dashed line) or 600 mV (yellow solid line) using PME with tinfoil boundary conditions (*ε′* = ∞). Red and purple lines: PMFs with the same external electric field as used for the 600 mV simulation but using non-conducting boundary conditions with permittivities *ε′* =1 (purple) or *ε′* = 80 (red).

The consequence of using vacuum boundary conditions instead of tinfoil boundary conditions is further illustrated from the distributions of water dipoles along open pores (Fig. S1). We find that external fields with vacuum boundary conditions (ε^’^ = 1) yield a dipole distribution that is symmetric with respect to the membrane center as requested for a nearly vanishing macroscopic dipole P, similar to the case without electric field. Consequently, neither the water in bulk nor the water in the pore are polarized, in sharp contrast to simulations with tinfoil boundary conditions. These data corroborate the observation from the PMFs that non-tinfoil boundary conditions suppress or even nearly remove effects from external electric fields. Hence, physically realistic simulations with external fields strictly require tinfoil boundary conditions.

## Discussion and Conclusions

The free energy landscape of membrane pore formation is highly sensitive with respect to transmembrane potentials, as evident from the modulations of pore free energies by tens to hundreds of kilojoules per mole by moderate potentials in the hundreds of millivolt range (Fig. 3). Hence, the equivalence of the PMFs obtained either with external electric fields or with charge imbalance suggests that the two methods impose equivalent electrostatic environments on the membrane. We believe that free energy calculations are a rigorous approach for excluding or for quantifying simulation artifacts. Free energy calculations may reveal moderate changes of thermodynamic driving forces which may not be apparent from the analysis of simulation structures alone. An example for such an artifact would be the free energy change of the open pore upon replacing tinfoil boundary conditions (Fig. 6, yellow, *ε′* = ∞) with water boundary conditions (Fig. 6, red, *ε′* = 80); this artifact may not be detected by merely observing pore formation, the pore structures, or dipole orientations (see Fig. S1). In a similar spirit, artifacts in non-neutral simulation systems with PME electrostatics have been quantified with free energy calculations^56^ and may be corrected during free energy calculations of ligated binding. ^57^

Our simulations together with previous results^12,22,34^ suggest that, both, external electric fields and charge imbalance provide valid and useful setups for simulating transmembrane potentials, given that tinfoil boundary conditions are applied. However, upon simulating constant potentials in the presence of conformational transitions that modulate the membrane capacitance, the charge imbalance method is technically more challenging since it requires updating of the charges. Since changes of membrane capacitance may only be estimated by geometric considerations, while updating of the charges may induce further alterations of membrane structure and capacitance, an iterative scheme for updating the charges may be required to reach a pre-selected potential. In contrast, imposing a pre-selected potential *V_m_* with external electric fields is simple since the external field *E_z_* is related to the transmembrane potentials via *V_m_* = *E_z_L_z_*, where *L_z_* is the box length in the direction of the electric field.

Membrane electroporation has been simulated extensively under non-equilibrium conditions by applying large transmembrane potentials of several volts.^7,8,10–12,23,24^ However, owing to the lack of good reaction coordinates of pore formation until recently, the free energy landscape of electroporation is not well understood. Such understanding will be critical for predicting the influence of different transmembrane potentials or of the lipid and protein content on the kinetics of pore opening and closure and, thereby, on the design of electric field pulse sequences with desired effects on cellular membranes. We anticipate that the present study lays the ground for deriving quantitative understanding of electroporation.

## Acknowledgement

We thank Bert de Groot for insightful discussions. This study was supported by Deutsche Forschungsgemeinschaft (DFG, German Research Foundation; grants SFB 803/A12 and SFB 1027/B7).

## TOC Graphic

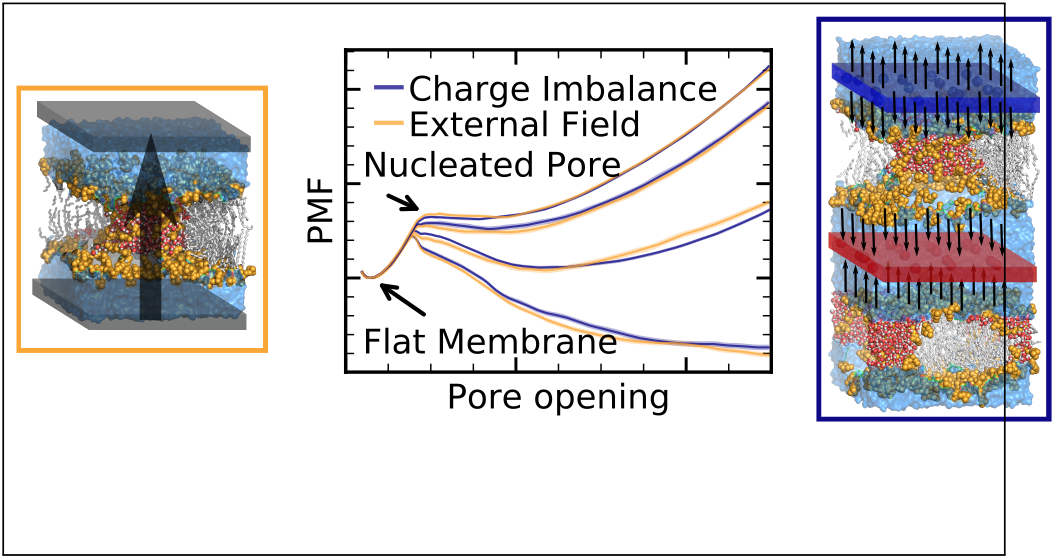

## References

(1) Allen, T. W.; Andersen, O. S.; Roux, B. Energetics of Ion Conduction through the Gramicidin Channel. PNAS 2004, 101, 117–122.

(2) Köpfer, D. A.; Song, C.; Gruene, T.; Sheldrick, G. M.; Zachariae, U.; de Groot, B. L. Ion Permeation in K^+^ Channels Occurs by Direct Coulomb Knock-On. Science 2014, 346, 352–355.

(3) Aksimentiev, A.; Schulten, K. Imaging α-Hemolysin with Molecular Dynamics: Ionic Conductance, Osmotic Permeability, and the Electrostatic Potential Map. Biophysical Journal 2005, 88, 3745–3761.

(4) Hub, J. S.; Aponte-Santamaría, C.; Grubmüller, H.; de Groot, B. L. Voltage-Regulated Water Flux through Aquaporin Channels in Silico. Biophys. J. 2010, 99, L97–L99.

(5) Jensen, M. Ø.; Jogini, V.; Borhani, D. W.; Leffler, A. E.; Dror, R. O.; Shaw, D. E. Mechanism of Voltage Gating in Potassium Channels. Science 2012, 336, 229–233.

(6) Vargas, E.; Yarov-Yarovoy, V.; Khalili-Araghi, F.; Catterall, W. A.; Klein, M. L.; Tarek, M.; Lindahl, E.; Schulten, K.; Perozo, E.; Bezanilla, F.; Roux, B. An Emerging Consensus on Voltage-Dependent Gating from Computational Modeling and Molecular Dynamics Simulations. Journal of General Physiology 2012, 140, 587–594.

(7) Tarek, M. Membrane Electroporation: A Molecular Dynamics Simulation. Biophys. J. 2005, 88, 4045–4053.

(8) Tieleman, D. P. The Molecular Basis of Electroporation. BMC Biochem 2004, 5, 1.

(9) Gurtovenko, A. A.; Vattulainen, I. Pore Formation Coupled to Ion Transport through Lipid Membranes as Induced by Transmembrane Ionic Charge Imbalance: Atomistic Molecular Dynamics Study. J Am Chem Soc 2005, 127, 17570–17571.

(10) Böckmann, R. A.; De Groot, B. L.; Kakorin, S.; Neumann, E.; Grubmüller, H. Kinetics, Statistics, and Energetics of Lipid Membrane Electroporation Studied by Molecular Dynamics Simulations. Biophys. J. 2008, 95, 1837–1850.

(11) Ziegler, M. J.; Vernier, P. T. Interface Water Dynamics and Porating Electric Fields for Phospholipid Bilayers. J. Phys. Chem. B 2008, 112, 13588–13596.

(12) Delemotte, L.; Tarek, M. Molecular Dynamics Simulations of Lipid Membrane Electroporation. J Membrane Biol 2012, 245, 531–543.

(13) Abidor, I.; Arakelyan, V.; Chernomordik, L.; Chizmadzhev, Y. A.; Pastushenko, V.; Tarasevich, M. Electric Breakdown of Bilayer Lipid Membranes I. The Main Experimental Facts and Their Qualitative Discussion. Bioelectrochem. Bioenerg. 1979, 6, 37–51.

(14) Benz, R.; Beckers, F.; Zimmermann, U. Reversible Electrical Breakdown of Lipid Bilayer Membranes: A Charge-Pulse Relaxation Study. J. Membrain Biol. 1979, 48, 181–204.

(15) Kotnik, T.; Rems, L.; Tarek, M.; Miklavčič, D. Membrane Electroporation and Elec-tropermeabilization: Mechanisms and Models. Annu. Rev. Biophys. 2019, 48, 63–91.

(16) Lambricht, L.; Lopes, A.; Kos, S.; Sersa, G.; Préat, V.; Vandermeulen, G. Clinical Potential of Electroporation for Gene Therapy and DNA Vaccine Delivery. Expert Opinion on Drug Delivery 2016, 13, 295–310.

(17) Aycock, K. N.; Davalos, R. V. Irreversible Electroporation: Background, Theory, and Review of Recent Developments in Clinical Oncology. Bioelectricity 2019, 1, 214–234.

(18) Geboers, B.; Scheffer, H. J.; Graybill, P. M.; Ruarus, A. H.; Nieuwenhuizen, S.; Puijk, R. S.; van den Tol, P. M.; Davalos, R. V.; Rubinsky, B.; de Gruijl, T. D.; Miklavčič, D.; Meijerink, M. R. High-Voltage Electrical Pulses in Oncology: Irreversible Electroporation, Electrochemotherapy, Gene Electrotransfer, Electrofusion, and Electroimmunotherapy. Radiology 2020, 295, 254–272.

(19) Zhong, Q.; Moore, P. B.; Newns, D. M.; Klein, M. L. Molecular Dynamics Study of the LS3 Voltage-Gated Ion Channel. FEBS Letters 1998, 427, 267–270.

(20) Tieleman, D.; Berendsen, H.; Sansom, M. Voltage-Dependent Insertion of Alamethicin at Phospholipid/Water and Octane/Water Interfaces. Biophysical Journal 2001, 80, 331–346.

(21) Roux, B. The Membrane Potential and Its Representation by a Constant Electric Field in Computer Simulations. Biophysical Journal 2008, 95, 4205–4216.

(22) Gumbart, J.; Khalili-Araghi, F.; Sotomayor, M.; Roux, B. Constant Electric Field Simulations of the Membrane Potential Illustrated with Simple Systems. Biochimica et Biophysica Acta (BBA) - Biomembranes 2012, 1818, 294–302.

(23) Piggot, T. J.; Holdbrook, D. A.; Khalid, S. Electroporation of the E. Coli and S. Aureus Membranes: Molecular Dynamics Simulations of Complex Bacterial Membranes. J. Phys. Chem. B 2011, 115, 13381–13388.

(24) Fernández, M. L.; Marshall, G.; Sagués, F.; Reigada, R. Structural and Kinetic Molecular Dynamics Study of Electroporation in Cholesterol-Containing Bilayers. J. Phys. Chem. B 2010, 114, 6855–6865.

(25) Sachs, J. N.; Crozier, P. S.; Woolf, T. B. Atomistic Simulations of Biologically Realistic Transmembrane Potential Gradients. J Chem Phys 2004, 121, 10847–10851.

(26) Denning, E. J.; Crozier, P. S.; Sachs, J. N.; Woolf, T. B. From the Gating Charge Response to Pore Domain Movement: Initial Motions of Kv1.2 Dynamics under Physiological Voltage Changes. Mol Membr Biol 2009, 26, 397–421.

(27) Kutzner, C.; Grubmüller, H.; De Groot, B. L.; Zachariae, U. Computational Electrophysiology: The Molecular Dynamics of Ion Channel Permeation and Selectivity in Atomistic Detail. Biophys. J. 2011, 101, 809–817.

(28) Ewald, P. The Calculation of Optical and Electrostatic Grid Potential. Ann. Phys. 1921, 64, 253–287.

(29) Darden, T.; York, D.; Pedersen, L. Particle Mesh Ewald: An N$\cdot$log(N) Method for Ewald Sums in Large Systems. J. Chem. Phys. 1993, 98, 10089–10092.

(30) Essmann, U.; Perera, L.; Berkowitz, M. L.; Darden, T.; Lee, H.; Pedersen, L. G. A Smooth Particle Mesh Ewald Potential. J. Chem. Phys. 1995, 103, 8577–8592.

(31) Raiteri, P.; Kraus, P.; Gale, J. D. Molecular Dynamics Simulations of Liquid–Liquid Interfaces in an Electric Field: The Water–1,2-Dichloroethane Interface. J. Chem. Phys. 2020, 153, 164714.

(32) Gurtovenko, A. A.; Anwar, J.; Vattulainen, I. Defect-Mediated Trafficking across Cell Membranes: Insights from in Silico Modeling. Chem. Rev. 2010, 110, 6077–6103.

(33) Kopec, W.; Gapsys, V. Periodic Boundaries in Molecular Dynamics Simulations: Why Do We Need Salt?; Preprint, 2022.

(34) Melcr, J.; Bonhenry, D.; Timr, Š.; Jungwirth, P. Transmembrane Potential Modeling: Comparison between Methods of Constant Electric Field and Ion Imbalance. J. Chem. Theory Comput. 2016, 12, 2418–2425.

(35) Hub, J. S. Joint Reaction Coordinate for Computing the Free-Energy Landscape of Pore Nucleation and Pore Expansion in Lipid Membranes. J. Chem. Theory Comput. 2021, 17, 1229–1239.

(36) Knight, C. J.; Hub, J. S. MemGen: A General Web Server for the Setup of Lipid Membrane Simulation Systems. Bioinformatics 2015, 31, 2897–2899.

(37) Berger, O.; Edholm, O.; Jähnig, F. Molecular Dynamics Simulations of a Fluid Bilayer of Dipalmitoylphosphatidylcholine at Full Hydration, Constant Pressure, and Constant Temperature. Biophys. J. 1997, 72, 2002–2013.

(38) Berendsen, H. J. C.; Postma, J. P. M.; DiNola, A.; Haak, J. R. Molecular Dynamics with Coupling to an External Bath. J. Chem. Phys. 1984, 81, 3684–3690.

(39) Abraham, M. J.; Murtola, T.; Schulz, R.; Páll, S.; Smith, J. C.; Hess, B.; Lindahl, E. GROMACS: High Performance Molecular Simulations through Multi-Level Parallelism from Laptops to Supercomputers. SoftwareX 2015, 1, 19–25.

(40) Bussi, G.; Donadio, D.; Parrinello, M. Canonical Sampling through Velocity Rescaling. J. Chem. Phys. 2007, 126, 014101.

(41) Hub, J. S.; Awasthi, N. Probing a Continuous Polar Defect: A Reaction Coordinate for Pore Formation in Lipid Membranes. J. Chem. Theory Comput. 2017, 13, 2352–2366.

(42) Awasthi, N.; Hub, J. S. Biomembrane Simulations. Computational Studies of Biological Membranes; CRC Press, Taylor & Francis Group, 2019.

(43) Ting, C. L.; Awasthi, N.; Müller, M.; Hub, J. S. Metastable Prepores in Tension-Free Lipid Bilayers. Phys. Rev. Lett. 2018, 120.

(44) Verbeek, S. F.; Awasthi, N.; Teiwes, N. K.; Mey, I.; Hub, J. S.; Janshoff, A. How Arginine Derivatives Alter the Stability of Lipid Membranes: Dissecting the Roles of Side Chains, Backbone and Termini. Eur Biophys J 2021, 50, 127–142.

(45) Poojari, C. S.; Scherer, K. C.; Hub, J. S. Free Energies of Membrane Stalk Formation from a Lipidomics Perspective. Nat Commun 2021, 12, 6594.

(46) Hub, J. S.; de Groot, B. L.; van der Spoel, D. G_wham–A Free Weighted Histogram Analysis Implementation Including Robust Error and Autocorrelation Estimates. J. Chem. Theory Comput. 2010, 6, 3713–3720.

(47) Gunsteren, W. F. V.; Berendsen, H. J. C. Gromos Manual; BIOMOS, Biomolecular Software, Laboratory of Physical Chemistry, University of Groningen, The Netherlands, 1987.

(48) Tieleman, D. P.; Leontiadou, H.; Mark, A. E.; Marrink, S.-J. Simulation of Pore Formation in Lipid Bilayers by Mechanical Stress and Electric Fields. J. Am. Chem. Soc. 2003, 125, 6382–6383.

(49) Tolpekina, T.; Den Otter, W.; Briels, W. Nucleation Free Energy of Pore Formation in an Amphiphilic Bilayer Studied by Molecular Dynamics Simulations. J. Chem. Phys. 2004, 121, 12060–12066.

(50) Bennett, W. D.; Sapay, N.; Tieleman, D. P. Atomistic Simulations of Pore Formation and Closure in Lipid Bilayers. Biophys. J. 2014, 106, 210–219.

(51) Awasthi, N.; Hub, J. S. Simulations of Pore Formation in Lipid Membranes: Reaction Coordinates, Convergence, Hysteresis, and Finite-Size Effects. J. Chem. Theory Comput. 2016, 12, 3261–3269.

(52) Hu, Y.; Sinha, S. K.; Patel, S. Investigating Hydrophilic Pores in Model Lipid Bilayers Using Molecular Simulations: Correlating Bilayer Properties with Pore-Formation Thermodynamics. Langmuir 2015, 31, 6615–6631.

(53) Weaver, J. C.; Chizmadzhev, Y. A. Theory of Electroporation: A Review. Bioelectrochemistry and Bioenergetics 1996, 41, 135–160.

(54) de Leeuw, S.; Perram, J.; Smith, E. Simulation of Electrostatic Systems in Periodic Boundary Conditions. I. Lattice Sums and Dielectric Constants. Proc. R. Soc. Lond. A 1980, 373, 27–56.

(55) Deserno, M.; Holm, C. How to Mesh up Ewald Sums. I. A Theoretical and Numerical Comparison of Various Particle Mesh Routines. J. Chem. Phys. 1998, 109, 7678.

(56) Hub, J. S.; de Groot, B. L.; Grubmüller, H.; Groenhof, G. Quantifying Artifacts in Ewald Simulations of Inhomogeneous Systems with a Net Charge. J. Chem. Theory Comput. 2014, 10, 381–390.

(57) Öhlknecht, C.; Lier, B.; Petrov, D.; Fuchs, J.; Oostenbrink, C. Correcting Electrostatic Artifacts Due to Net-charge Changes in the Calculation of Ligand Binding Free Energies. J Comput Chem 2020, 41, 986–999.

