## Supplemental PDF for "Equivalence of charge imbalance and external electric fields during free energy calculations of membrane electroporation"

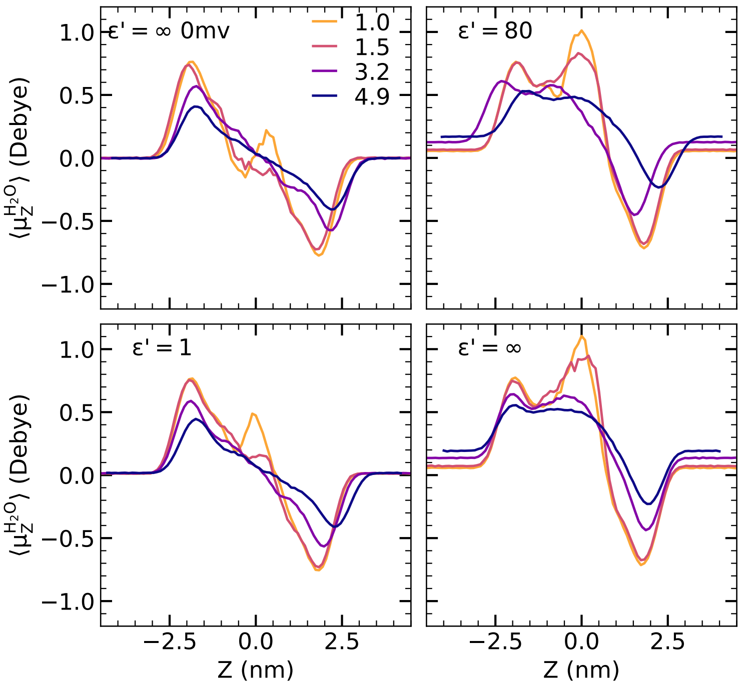


Figure S1: On the effect of PME boundary conditions on the polarization of water. (A) For reference, profiles of the average dipole per water molecule projected on the $z$ axis, $\langle \mu_z^{H_2O} \rangle$, for simulations with zero transmembrane potential and using tinfoil boundary conditions ($\varepsilon' = \infty$). Profiles are shown for defects of increasing size as given by values of the coordinate $\xi_p = 1.0$, 1.5, 3.2, or 4.9 in shades from yellow to blue. (B–D) Corresponding profiles using an external electric field that, using tinfoil boundary conditions, would yield a transmembrane potential of 600 mV. Profiles are shown for PME boundaries with (B) $\varepsilon' = 80$, (C) $\varepsilon' = 1$, and (D) $\varepsilon' = \infty$. Using $\varepsilon' = 1$ suppresses the formation of a macroscopic dipole as evident from dipole distributions similar to the case without external field (compare C with A). In contrast, using $\varepsilon' = 80$ leads to similar dipole distributions as compared to tinfoil boundary conditions (compare B with D), despite the fact that $\varepsilon' = 80$ reduced the electric-field effect on the free energy of the open pore by $\sim$50 kJ/mol. Hence, the dipole distribution alone is an insufficient indicator for a correct implementation of electric fields.
